# Biotic challenges in the city: Dietary restrictions and body fat content of female sexuals in urban ant populations

**DOI:** 10.1101/2024.04.01.587520

**Authors:** Gema Trigos-Peral, Magdalena Witek, Enikő Csata, Paulina Chudzik, Jürgen Heinze

## Abstract

Urban habitats represent an important challenge for many organisms. Besides the abiotic changes, urban habitats are also characterized by changes in the biotic conditions, such as a more uniform species composition and declining population sizes. For urban ants this can result in dietary shortages. In our study, we tested whether urban ant colonies might suffer from dietary restrictions by carrying out a common garden experiment in which ant colonies from urban and rural habitats were exposed to high carbohydrate, protein, and fat / protein diets. We also investigated the body fat content of individuals from both habitat types. Our findings suggest a lower availability of high-quality carbohydrates in urban areas. Additionally, while not statistically significant, rural colonies exhibited a tendency to consume greater quantities of proteins and fat compared to urban colonies. This trend was in line with a higher body fat content observed in female sexuals (gynes) from rural colonies. These results might indicate the outcome of an evolutionary feedback process in which ant colonies adapt to nutritional constraints in urban environments. They achieve this by minimizing the investment in gynes, which might require fewer reserves for survival during colony foundation due to reduced competition for nesting sites within urban green spaces.

## Introduction

During the last decades, urban ecology has received an increasing interest in the scientific community. Several studies have demonstrated how the anthropogenic transformation of the habitat has caused numerous behavioural and physiological changes in organisms (Ditchkoff et al., 2006; Calfapietra et al., 2015; Brans et al., 2018; Łopucki et al., 2021; Jacquier et al., 2023; Trigos-Peral et al., 2024; among others). The best-known consequences of urbanization are the biotic homogenization of the habitat and the establishment of invasive species (Vilà et al., 2011; Trigos-Peral et al., 2020, 2021; Brassard et al., 2021). Both affect species composition and habitat-niche partitioning, thereby triggering additional alterations in the local ecological networks. For example, the decrease in the availability or quality of food sources can disrupt fragile food webs (Shrewsbury & Raupp, 2006; Qu et al. 2022), influence food size preferences (Chen & Neoh 2023) and threaten the survival of species.

Low nutritional quantity and quality negatively affect growth, behavioural performance, reproduction, immunological defence, and lifespan (i.e., Spann & Schumann, 2009; McWilliams, 2011; Simpson & Raubenheimer, 2012; Nwaogu et al., 2020; Shik & Dussutour, 2020; Abellán et al., 2023) and thus endanger the survival of both species of plants and animals. Scarcity and low quality of nutrition particularly challenge organisms with low mobility, in which the chances of moving to new areas to find more suitable food sources are limited. For example, most ants are central-place foragers and their foraging efficiency is influenced by the distance between their stationary nest and the food source (Devigne & Detrain, 2006). Though many ant species have adapted to the ecological conditions in urban areas (Brassard et al., 2021), the potential nutritional deficiencies in this habitat type might nevertheless influence individual survival or brood development and consequently affect colony growth.

Ants obtain essential micronutrients, such as vitamins, salts, amino acids, or lipids, by consuming macronutrients from plant and animal sources. They gain proteins, fat, and salts by hunting or feeding on cadavers and in many species, carbohydrates and essential amino acids are provided in secretions from aphids (honeydew) (Csata et al., 2020). Honeydew is an easily accessible nutritional source due to the ubiquitous and cosmopolitan distribution of aphids (Alyokhin et al., 2022). Several aphid species appear to benefit from urban habitats, in which the number of their predators may be substantially reduced (Korányi et al., 2020). Nevertheless, honeydew composition heavily relies on the host plant species and its nutritional state (Fischer et al., 2005; Leroy et al., 2011). It is therefore possible that honeydew from urban aphids is of lower quality. The combined effects of a decline in ground-dwelling arthropods (Vergnes et al., 2013) and the potential impoverishment of aphid honeydew may jeopardize the variety and quality of the diet of ants.

In addition to carbohydrates and proteins, lipids are also important macronutrients (Blüthgen & Feldhaar, 2009), which influence the growth and survival of ant colonies. Ants can obtain fatty acids either directly from their diet or synthesise them from carbohydrates (Heath & Rock, 2002). Fat stored in the fat body of insects is crucial in numerous physiological processes. For example, it increases resistance to starvation (Dussutour et al., 2016), stimulates egg development (Legaspi & Legaspi, 1998; Legaspi & O’Neil, 1994), and affects the division of reproductive labour in queenless ants (Bernadou et al., 2020).

We therefore hypothesize that urban habitats represent an important challenge to the survival of ant colonies due to the potentially reduced quantity and quality of available natural food sources (i.e., Kaspari et al., 2008; Bujan & Kaspari, 2017; Kaspari et al., 2019). In a common garden study, we compared the foraging preferences of the ant *Lasius niger* from rural areas and urban parks to determine whether there is a nutritional shortage in the city and how it impacts ant colonies. Specifically, we investigated i) whether urban and rural colonies differ in their preference for three types of diet with different carbohydrate, protein, and fat content, and ii) if workers and female sexuals (gynes) from the two types of habitat differ in body fat content.

## Material and Methods

### Food source choice and consumption

The study was carried out in July 2021 using colonies of *Lasius niger* collected in three urban parks located in the city centre of Warsaw, Poland, and three rural areas (free of human influence) in the surroundings (Supplementary table 1). We collected three colony fragments in each of three urban and three rural areas (total 18 colony fragments). Each fragment consisted of a minimum of 500 workers, a few hundred larvae, and a few dozen pupae of female sexuals (gynes) and workers (differenced by the size and shape).

Immediately after collection, from each fragment we set up two experimental colonies consisting of 150 workers each. Brood was not included since it would have influenced both the food selection and the amount consumed depending on their developmental status, which could not easily be standardized (Dussutour & Simpson, 2008). The workers were placed in small plastic containers (18 cm × 12 cm × 6 cm) with a humid piece of sponge covered by a dark plastic plate serving as nest. Each container was connected to three different foraging arenas (10 cm × 6.5 cm × 5 cm) via silicone tubes. The 36 experimental colonies were deprived of food for one day, while the ants remaining in the18 original fragments were provided with sugar-water and mealworms *ad libitum* until they were used in the later experiments.

After starving the experimental colonies for one day, we started the nutrition choice experiment by placing into each foraging arena a feeder (a plastic plate of 3 cm diameter) with a different type of food: carbohydrates, protein, and fat / protein. The carbohydrate source was a saturated sucrose solution (1 M), protein was canned tuna packed in its own juice, and the fat / protein source was canned tuna in soybean oil (Supplementary table 2). To avoid differences in food retrieval caused by different food consistency, both protein and fat / protein were blended using mineral water (same that the one used for the sucrose solution) to achieve a liquid consistency similar to the carbohydrate solution. After placing the food sources, we opened the access to the arena and recorded the feeding activity of ants for half an hour using a video camera (Sony HDR-AS20 Full HD). Food sources were placed randomly across the observations.

One week later we repeated the experiment to determine whether food preferences are consistent or change after *ad libitum* feeding in the laboratory. The 36 experimental colonies were maintained at room temperature (approximately 24 °C) under a natural light regimen and fed *ad libitum* by each day randomly providing new feeders with fresh prepared carbohydrates, protein, and fat / protein diets. After one week, we starved the colonies again for one day and then repeated the food preference experiment.

To estimate the food choice by the ants, we analysed the videos and counted the total number of ants present in each foraging arena every five minutes. To determine how much food was consumed by the ant workers, we weighed the amount of each type of food placed in the feeders before and after each observation using a laboratory balance (Ohaus Scout SKX123; error: ± 1 mg).

### Body fat content and gyne size

To test if body fat content is higher in gynes of rural colonies due to the presumed richer nutritional availability, we collected five gynes and ten workers from five nests each located in the three urban parks and rural areas that had been sampled for the food choice experiment (Supplementary table 1). Samples were collected shortly before the massive nuptial flight of *L. niger*, which occur in Poland in mid-July (Van der Have et al. 2011 and pers. observations). We therefore can assume that all gynes collected for this experiment were produced approximately at the same time, since gynes that had been produced earlier should have already abandoned the nest during previous small nuptial flights. Each collected individual was placed in an individually labelled 2 ml vial and frozen at −25 °C. To determine whether the differences in fat content reflected differences in the food consumed in the two different habitat types or differences in the genetic background of the two populations, we also measured body fat content in gynes and workers born and reared in the laboratory in the colonies collected one month before for the previous experiment and equally fed *ad libitum* with identical food (honey water and tenebrids/*Drosophila*).

Body fat content was measured following Bernadou et al. (2015). Individuals were dried at 60 °C for five days and weighed individually to the nearest 0.0001 mg with a microbalance (Radwag MYA 5.4Y) to measure their dry mass. Subsequently, fat was extracted by soaking each worker for two days in 2 ml petroleum ether (boiling range 40–60 °C, Merck, Darmstadt, Germany) at room temperature. After two days, the workers were transferred into new vials, the petroleum ether was renewed, and each worker was kept in petroleum ether again for two days. Afterwards, the ants were dried at 60 °C for six days and weighed again to determine their lean mass. The percentage of fat was calculated according to the standard equation: (dry mass − lean mass) * 100/dry mass.

## Statistical analyses

### Food source choice and consumption

We tested whether urban and rural colonies were differently attracted by the different types of food (carbohydrates, protein, or fat / protein) sources by carrying out a GLMM (zero inflated Poisson) in which the number of foragers observed in each foraging arena was used as the dependent variable. The three-way interaction of habitat type, type of food, and the round of the experiment was included as explanatory factors, colony identity, and round of observation as nested random factors.

We checked whether the total amount of food consumed differed between habitat types and repetitions by carrying out a LMM (Gamma distribution) in which the total amount of food consumed was used as explanatory variable and the interaction between habitat type and round of the experiment as explanatory factors. We also tested for differences in the actual quantity of each type of food consumed by the colonies from each habitat type and across rounds with a LMM approach in which the *log+1* transformed quantity of carbohydrates, protein, fat / protein consumed was used as response variable, and the interaction between habitat type, type of food, and round of the experiment were used as the explanatory factors. In all LMMs, colony identity was included as random factor. Finally, paired t-tests were carried out on the consumption of each nutrient per each population to compare differences between the first and second round of observations.

### Body fat content and gyne size

To test for differences in body fat content between ants from urban and rural habitats, we performed a LMM with fat content as response variable and the interaction of the habitat type, caste, and origin (field or laboratory) as explanatory factors. The origin of the individuals was included in the model to avoid an influence of genetic background. Finally, since larger gynes can potentially accumulate larger quantities of body fat, we tested for size differences in gynes between habitat types using dry mass as a proxy of individual size. Comparisons were performed using a LMM approach, in which individual dry mass was introduced as dependent variable, the interaction between habitat type and origin of the individual (field or laboratory) as explanatory factors, and colony identity was used as random factor.

Statistical analyses were performed using R v4.1.2 (R Core Team, 2021) and RStudio (Posit Team, 2023). Linear mixed models (LMM) were performed by using the function *lmer* from the package *lme4* (Bates et al., 2013). Models were tested for overdispersion by using the function *testdispersion* of the package DHARMa (Hartig, 2022). When overdispersion was found in LMMs, the dependent variable was transformed using the most suitable transformation approach obtained using the function *bestNormalize* from the R-package bestNormalize (Peterson, 2021). Zero-inflated Poisson regression was performed by using the *zeroinfl* function of the pscl package (Jackman, 2020). The significance of the explanatory factors was determined by performing the Anova of the LMMs and GLMMs using the function *Anova* from the car package (Fox & Weisberg, 2019). To assess differences in urban-rural trends for the tested factors, we performed post-hoc pairwise comparisons of all models using the *emmeans* function of the emmeans package (Lenth, 2023) for GLMMs. For LMMs we used the *posthoc_Pairwise* function of the grafify package (Shenoy, 2021). Graphical representations were performed using the packages ggplot2 (Wickham, 2016), magick (Ooms, 2023), png (Urbanek, 2022) and ggpubr (Kassambara, 2022). Mean and error values were calculated using the Rmisc package (Hope, 2022).

## Results

### Food source choice and consumption

Our results show that the interest of workers in exploring the different arenas was significantly influenced by food type (χ = 41.64, p < 0.001), habitat type (z = 6.84, p < 0.001), the round of the experiment (χ = 15.06, p < 0.001), and the interactions between these factors (Table 1). Just after collection (round 1), workers of urban and rural colonies did not differ in their interest in carbohydrates (z = −0.09; p = 1). In contrast, rural colonies were significantly more foraging for protein and fat / protein diets (protein: z = 3.85, p = 0.006; fat / protein: z = 3.31, p = 0.043). One week later (round 2), rural colonies were less interested in carbohydrates (z = −5.21, p < 0.001) and more interested in the fat / protein diet (z = 4.83, p < 0.001) than urban colonies (Figure 1).

**Figure 1.**
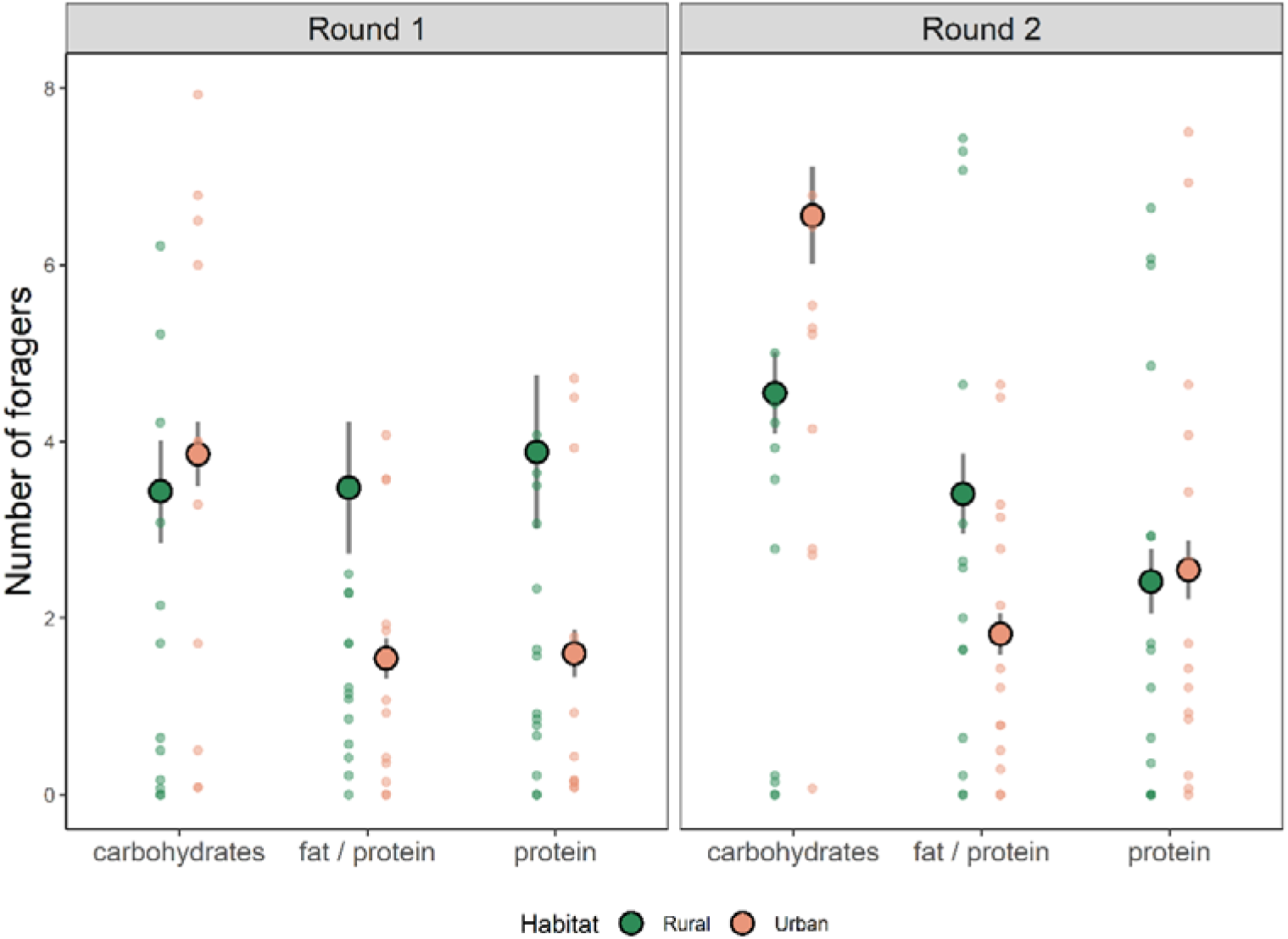
Mean and error bar plot showing the difference in the number of workers of the ant *Lasius niger* from rural and urban colonies present in the foraging arena containing different types of diet. The filled circles indicate the mean value of the consumption by the rural and urban colonies, the whiskers represent the standard deviation and the dots are the individual worker numbers for each colony.

**Table 1.** Influence of the study factors in the foraging interest of the workers in the food selection experiment.

| Number of workers attracted by each type of food |  |  |  |
| --- | --- | --- | --- |
| Factor | Df | $\chi^2$ | p-value |
| Type of food | 2 | 41.64 | < 0.001 |
| Habitat type | 1 | 6.84 | < 0.001 |
| Round of observation | 1 | 15.06 | < 0.001 |
| Type of food * Habitat type | 2 | 57.45 | < 0.001 |
| Type of food * Round of observation | 2 | 15.96 | < 0.001 |
| Habitat type * Round of observation | 1 | 26.84 | < 0.001 |
| Type of food * Habitat type * Round of observation | 2 | 6.72 | 0.035 |

When investigating the actual food intake, the total food consumed by the colonies decreased in the second round of the experiment (t = −2.19, p = 0.028; Supplementary figure 1). Colonies from urban and rural habitats consumed different quantities of each diet in the first round of observations. After collection, urban ants consumed more carbohydrates than rural colonies (t = 3.75, p < 0.001), but no clear differences were found in the consumption of protein (t = 0.75, p = 0.675) and fat / protein diets (t = 0.05, p = 0.971).

After one week of *ad libitum* feeding in the laboratory (round 2), no significant differences were found in food consumption between ants coming from different habitat types (carbohydrates: t = 0.07, p = 0.971; protein: t = 0.41, p = 0.848; fat / protein: t = 0.137, p = 0.971; Supplementary figure 2).

Finally, we observed a consistent decrease in the consumption of various types of nutrients the experimental colonies from both rural and urban populations between the first and second round of observations (Figure 2, Supplementary table 3).

**Figure 2.**
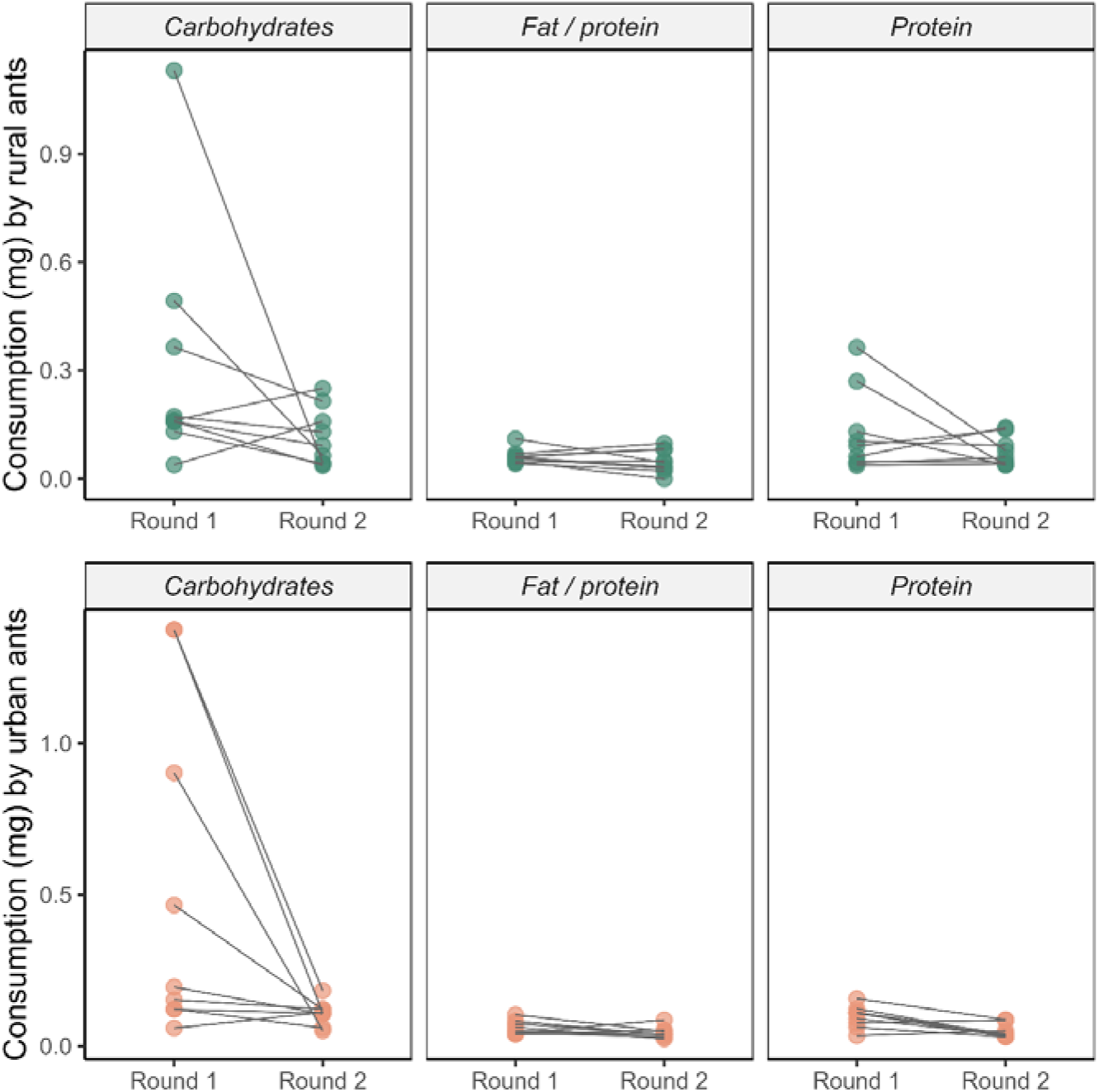
Paired dot plot showing the consumption of each diet (mg) by *Lasius niger* workers from urban and rural colonies during the two rounds of the experiment (one day after collection and one week after collection and feeding on a standard diet).

**Figure 3.**
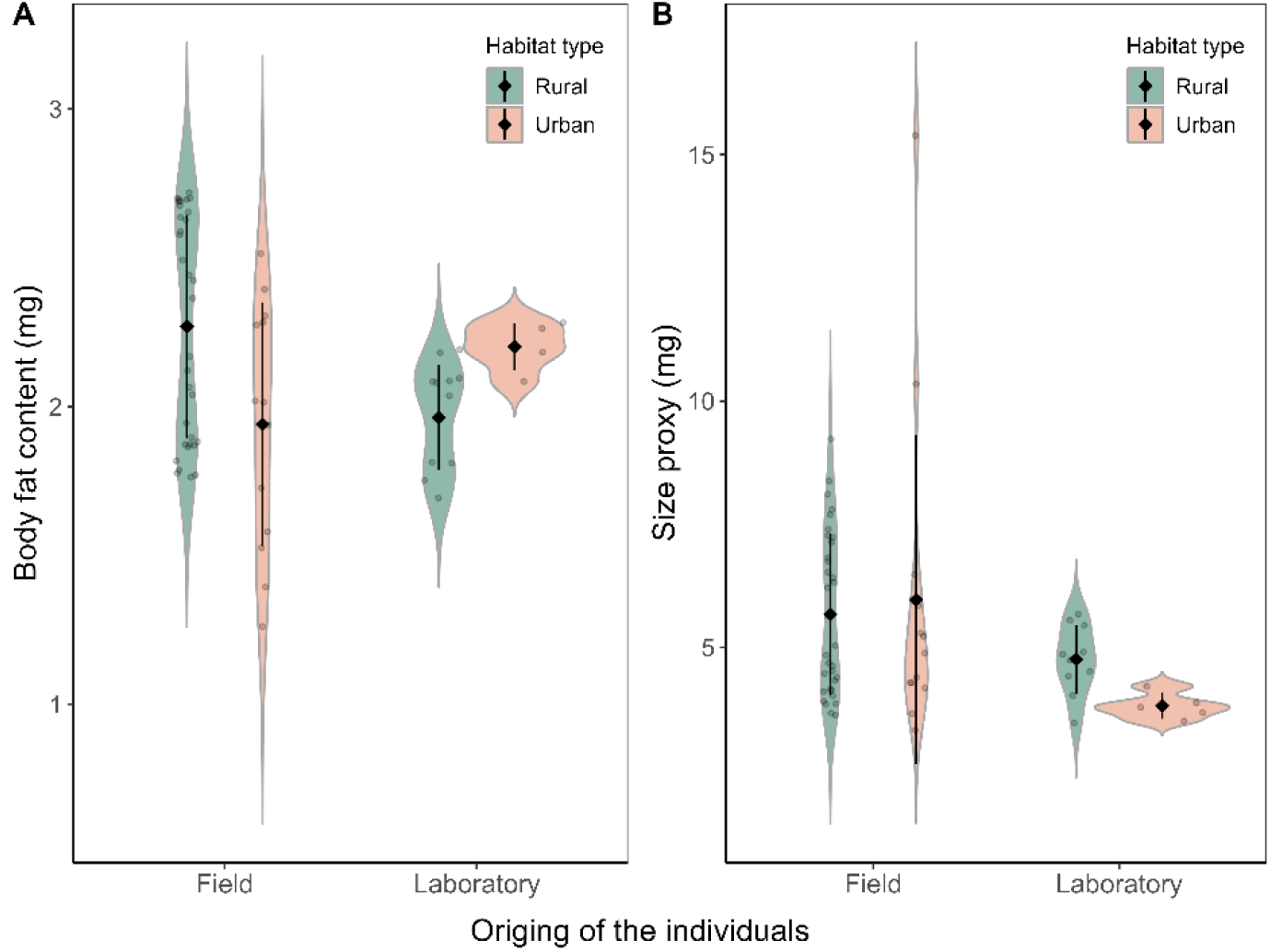
Violin plot and mean and error plot showing the body fat content and (in milligrams) size (using the dry mass as a proxy) of gynes of the ant *Lasius niger* reared under natural conditions in the field and under laboratory conditions. The black diamonds indicate the mean value of the consumption by the rural and urban colonies, and the whiskers represent the standard deviation.

### Body fat content and gynes size

Body fat content of gynes collected in the field differed between urban and rural habitats. Rural gynes had a significantly higher fat content than urban gynes (t = 3.52, p = 0.003), although laboratory-reared gynes from the two habitats did not differ (t = −1.51, p = 0.262). Workers from the two habitat types did not differ in body fat content, regardless of whether collected from the field (t = 0.78, p = 0.635) or reared in the laboratory (t = −1.76, p = 0.169). Finally, using the dry mass (mg) as a proxy of individual size, we found no significant difference between urban and rural gynes collected either in the field (t = −0.13, p = 0.895) or reared in the laboratory (t = −1.93, p = 0.072).

## Discussion

Workers of the ant *Lasius niger* from urban and rural colonies exhibited distinct preferences for three offered diets. Whereas the foraging activity in the arena containing carbohydrates did not differ between colonies from the two habitat types, workers from rural colonies showed significantly more interest in diets rich in protein or fat and protein compared to workers from urban ones.

Immediately after collection, workers from urban colonies showed a significantly higher intake of carbohydrates than workers from rural colonies, while no significant differences were found in the consumption of the other two diets. However, after one week of feeding on a standard diet in the laboratory, rural ants exhibited a significantly greater interest for the diet rich in fat / protein and less interest in carbohydrates, in contrast to urban ants. Ants from urban and rural habitats did not differ in the quantities they consumed from each diet.

Gynes from rural colonies were similar in size to those from urban colonies but had a significantly higher fat content. No differences in fat content were found when gynes from different habitat types were reared in the laboratory. Fat content was also identical in workers from urban and rural areas regardless of whether they were reared in the field or the laboratory.

We found a high general attraction of the ants for carbohydrates, which matches the well-known symbiotic association of *L. niger* with aphids (Czechowski et al., 2012; Seifert, 2018). The higher intake of carbohydrates in urban colonies immediately after collection suggests that carbohydrate sources, mainly honeydew, are less abundant and potentially of lower quality in urban habitats compared to sources in rural areas. Lower plant diversity and their growth in poor soil (Lescano et al., 2022) might negatively affect the honeydew production by aphids (Fisher et al., 2005) in urban areas, although it is worth noting that no specific studies have yet been carried out to test this assumption.

Contrary to our expectation that urban colonies would consume more protein and fat than rural colonies, we found that rural colonies were more interested in these diets, although differences were not significant. However, we attribute the lack of significance to the high concentration of nutrients in these diets, as found that ants the volume of food ingested by ants decreases with the increase of concentration in the diet (Dussutour & Simpson, 2008). Besides, we cannot exclude that recruitment during the initial observation round (one day after collection) still reflected natural factors, such as colony size and productivity (Mailleux et al., 2003). For example, colonies in rural areas might have been larger and more productive than urban colonies, explaining the unexpected, increased interest in protein and fat. While urban colonies can prolong their production period due to the elevated temperatures in urban environments (Trigos-Peral et al., 2024), rural colonies tend to have shorter reproduction periods and may need to optimize their productivity particularly during summer. As a result, rural colonies might have a greater demand for protein and fat to ensure an adequate brood development (Dussutour & Simpson, 2008; Rosumek et al., 2017).

Moreover, rural habitats present additional challenges in the search for proteins and fats. While greater biodiversity can theoretically provide a wider array of food sources, it also results in an increased number of neighbouring colonies competing for these resources. Penick et al. (2015) suggested that certain urban ant species can compensate the lack of natural food sources by feeding on human food waste. However, the observed consumption of leftover food varied with the ants’ origin (native or invasive), location (parks or traffic islands), and trophic level (predator or herbivore). For example, consumption of food waste was only evidenced in the omnivorous, exotic ants *Tetramorium* spp. and *Nylanderia flavipes* in traffic islands, while *Lasius cf. emarginatus* appeared to avoid human food waste (Penick et al., 2015).

After being provided with similar diets for one week in the laboratory, rural colonies not only again foraged more for fat / protein but also exhibited an increased consumption of this diet. Though this might suggest that rural ants suffer more from protein and fat shortage, we found that rural gynes had a higher body fat content than urban counterparts. Body fat content plays a critical role in the success of solitary colony foundation, as it significantly affects queen survival and egg production during the development of the first workers (Heinze & Tsuji, 1995; Helms & Kaspari, 2015; Negroni et al., 2021). Body fat content increases in gynes after emergence due to feeding (Boomsma & Isaaks, 1985), so the differences might in principle result from a later emergence of urban gynes. However, the annual activity starts earlier in urban ants than in rural ones (GT unpublished data), which should lead to opposite results (higher body fat content in urban ants). Furthermore, prior to the massive nuptial flight in July, there are a few smaller nuptial flights in which the earlier emerged gynes leave the nest. Notwithstanding, further studies are needed to determine whether the lower fat content observed in urban gynes might result from different timing of emergence or indeed indicates an evolutionary adaptation to dietary constraints in their habitat. Moreover, although workers from both habitat types did not differ in body fat content, the nutritional shortage may not be apparent in this caste throughout our experiment. For instance, although Rosumek et al. (2017) did not find differences in body fat content of ant workers when testing diets with low and high fat content, they found a change of their fatty acid profile.

To summarize, our research offers new insights into the potential dietary shortages in urban areas and its consequences on ants. Further studies with controlled diets and genetic approaches are needed to accurately determine the interplay between the availability of food sources, foraging behaviour, and reproductive success in ants. In addition, our findings highlight once more the significance of diet quality in ants (Csata et al., 2020; Wendt & Czaczkes, 2020) and emphasize the importance of meticulously selecting the diets for use in baiting assays.

## Data availability

The data used in this study are available through the link to figshare (https://figshare.com/s/20b4ecf6e7b571b9ca13) in the manuscript. Scripts and outputs are also available through the former link. Doi: 10.6084/m9.figshare.25513309

## Supplementary material

**Supplementary table 1.** Locations for the collection of the fragments of urban and rural colonies in Warsaw (Poland).

| Urban areas |  |
| --- | --- |
| Location | GPS coordinates |
| *Ogród Saski Park* | 52.240482, 21.008921 |
| *Morskie Oko Park* | 52.205943, 21.026827 |
| *Marshal Edward Rydz-Śmigły Park* | 52.225446, 21.033746 |
| Rural areas |  |
| Location | GPS coordinates |
| *Gmina Michałowice* | 52.159636, 20.873293 |
| *Klaudyn* | 52.272952, 20.861246 |
| *Warsaw Airport* | 52.146279, 20.987392 |

**Supplementary table 2.** Nutritional information in 100 gr of canned tuna in own juice (protein) and tuna in soybean oil (fat / protein) used in the experiments.

| Proteins (tuna in own juice) |  |
| --- | --- |
| Nutrient | In 100 gr of product |
| *Proteins* | 23 gr |
| *Fat* | 0.6 gr |
| *Salt* | 1.05 gr |
| *Carbohydrates* | < 0.1 gr |
| Fat / proteins (tuna in soybean oil) |  |
| Nutrient | In 100 g of product |
| *Proteins* | 23.5 gr |
| *Fat* | 9.5 gr |
| *Salt* | 1.05 gr |
| *Carbohydrates* | < 0.1 gr |

## Supplementary figures

**Supplementary figure 1.**
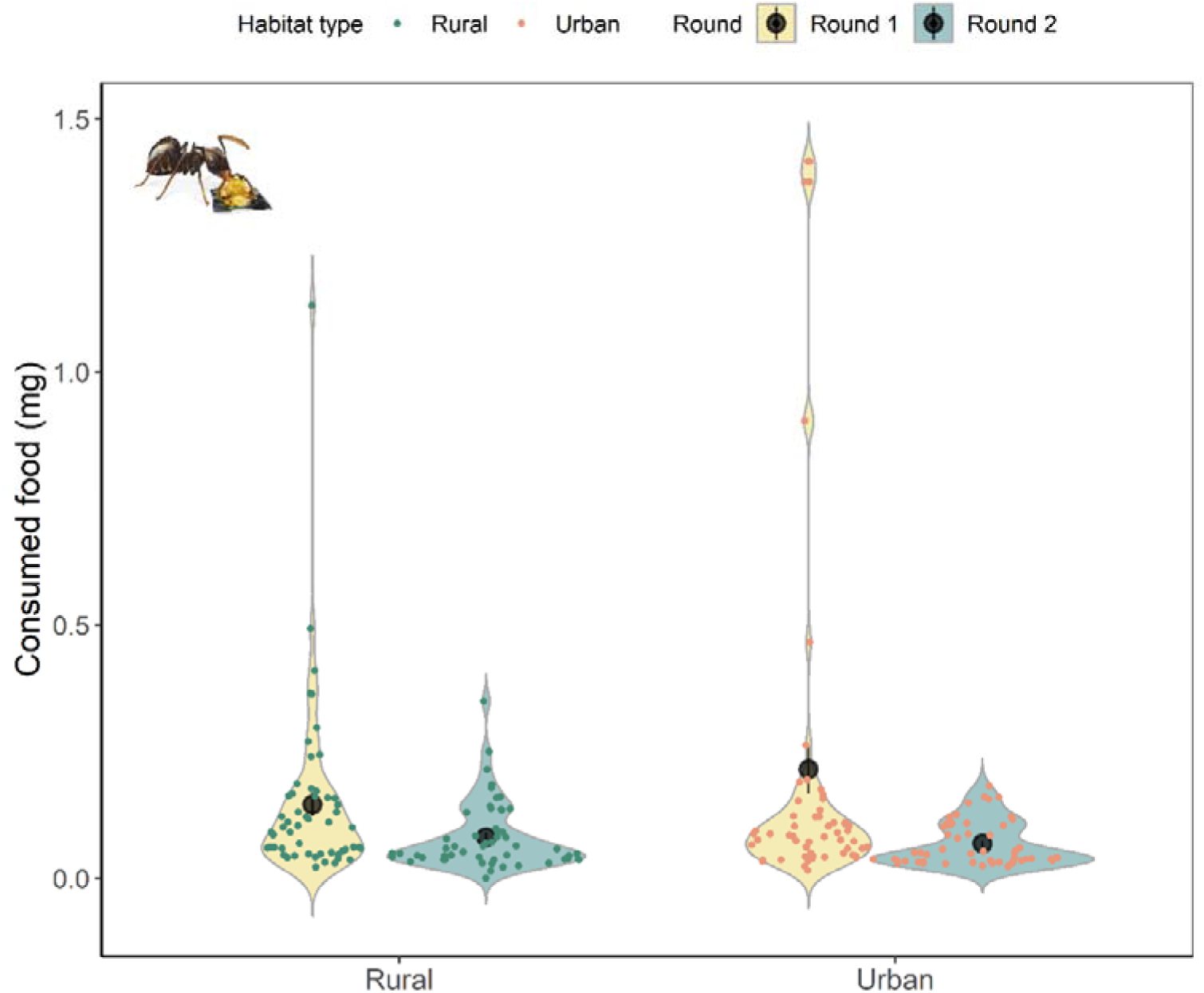
Violin plots showing the amount of food consumed (mg) by the colonies from urban and rural habitats during the first round of observations (one day after colonies collection) and the second round of observations (one week after collection and feeding at the laboratory under the same type of diets). The black circles show the average consumption per each population, whereas coloured dots represent the individual values for each urban (in pink) and rural (in green) experimental colony.

**Supplementary figure 2.**
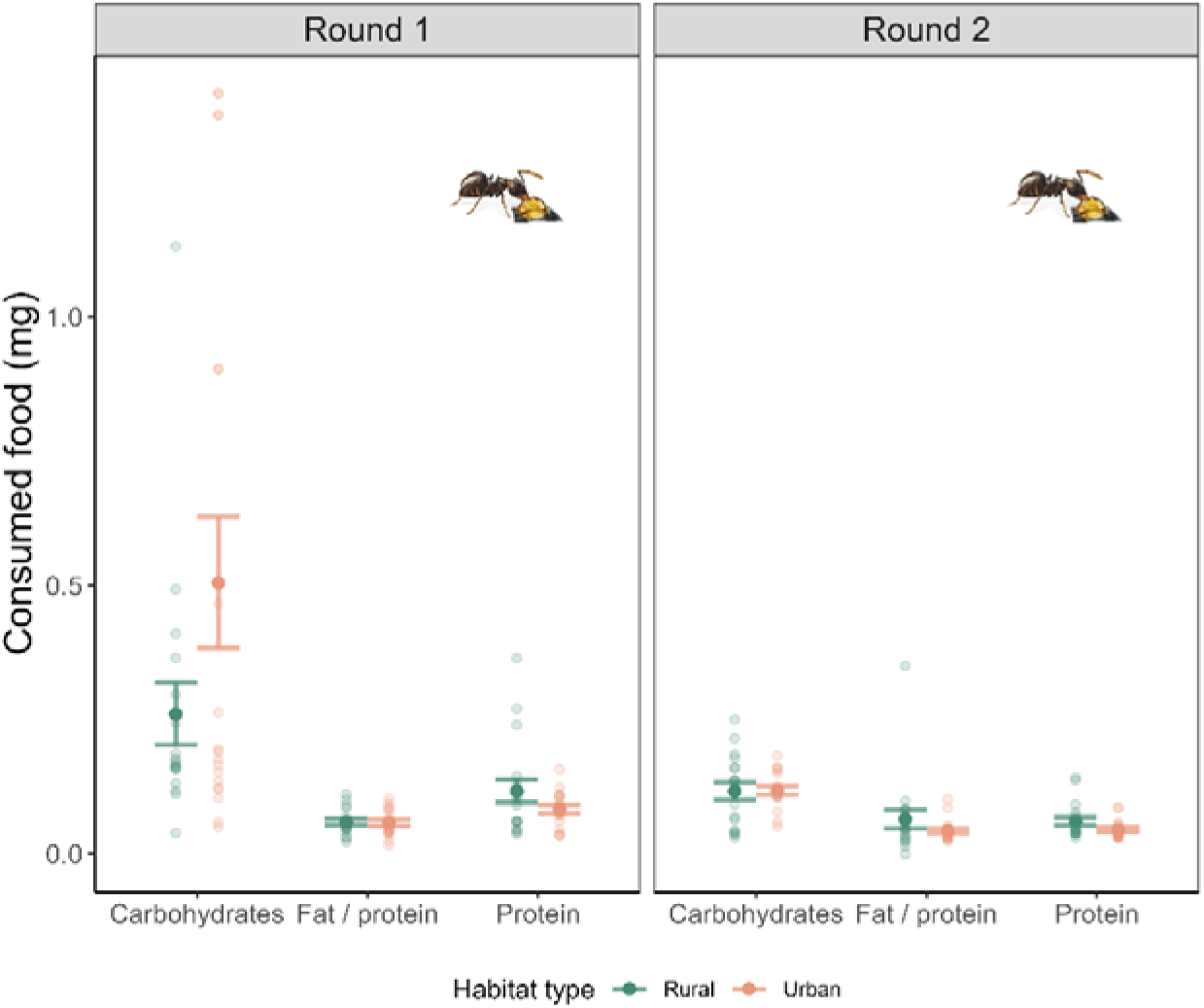
Mean and error bar plot showing the difference in the amount of food (mg) consumed by *Lasius niger* workers from urban and rural colonies during the two rounds of the experiment (one day after collection and one week after collection and feeding on a standard diet). The full circles indicate the means of the consumption by the rural and urban colonies, and the whiskers represent the standard deviation. Open circles represent the individual values per each experimental colony.

